# Load-dependent modulation of alpha oscillations during working memory encoding and retention in young and older adults

**DOI:** 10.1101/848127

**Authors:** Sabrina Sghirripa, Lynton Graetz, Ashley Merkin, Nigel C Rogasch, John G Semmler, Mitchell R Goldsworthy

## Abstract

Working memory (WM) is vulnerable to age-related decline, particularly under high loads. Visual alpha oscillations contribute to WM performance in younger adults, and although alpha decreases in power and frequency with age, it is unclear if alpha activity supports WM in older adults. We recorded electroencephalography (EEG) while 24 younger (aged 18-35 years) and 30 older (aged 50-86) adults performed a modified Sternberg task with varying load conditions. Older adults demonstrated slower reaction times at all loads, but there were no significant age differences in accuracy. Regardless of age, alpha power decreased, and alpha frequency increased with load during encoding, and the magnitude of alpha suppression during retention was larger at higher loads. While alpha power during retention was lower than fixation in older, but not younger adults, the relative change from fixation was not significantly different between age groups. Individual differences in alpha power did not predict performance for either age groups or at any WM loads. Future research should elaborate the functional significance of alpha power and frequency changes that accompany WM performance in cognitive ageing.

## 1 Introduction

Verbal working memory (WM), the ability to actively maintain and/or manipulate verbal information to guide immediate cognitive processing (Baddeley, 1992), is vulnerable to age-related decline. Compared to younger adults, healthy older adults are able to store fewer items in WM, are less able to manipulate those items (Fisk and Warr, 1996), and are more susceptible to interference from distracting information (Gazzaley and D’esposito, 2007). This age difference is particularly salient under high WM demands, with older adults demonstrating poorer performance with higher loads relative to younger adults (McEvoy et al., 2001; Wild-Wall et al., 2011). Despite this, the neural mechanisms underlying such age differences at varying WM loads are not well understood.

Advancing age is associated with progressive changes in the frequency and power of neural oscillations (Klass and Brenner, 1995; Klimesch, 1999). Alpha (∼8-12Hz) is perhaps the most affected frequency band in ageing, with alpha oscillations significantly lower in magnitude and slower in frequency in healthy older adults compared with younger adults at rest (Babiloni et al., 2006; Klimesch, 1997; Lindsley, 1939). As alpha oscillations in posterior brain regions are thought to support WM performance (Klimesch, 2012), age-related changes to alpha activity may underlie WM performance deficits in healthy older adults.

WM is typically divided into three stages: encoding, retention and retrieval (Baddeley, 1992). Most of the research in this area has focused on the retention period, with a large body of evidence suggesting that alpha is modulated during this stage, though the location, direction and magnitude of this change depends on the type of task. Using modified Sternberg tasks, it has been reliably shown that alpha power increases in visual brain areas during retention, particularly under higher WM loads (Jensen et al., 2002; Meltzer et al., 2008; Proskovec et al., 2019). The predominant interpretation of this finding is that alpha activity reflects a suppression of sensory input from the visual stream to prevent disruption to WM maintenance occurring in frontal brain regions (Jensen and Mazaheri, 2010). In lateralised WM tasks where subjects attend to and memorise the information in one hemifield, and ignore the other, parieto-occipital alpha power decreases in the task-relevant, and increases in the task-irrelevant hemisphere (Sauseng et al., 2009). Finally, alpha suppression with increasing WM load in parieto-occipital sites has been reported in n-back style paradigms (Gevins et al., 1997; Krause et al., 2000; Pesonen et al., 2007; Stipacek et al., 2003) and delayed match-to-sample tasks (Fukuda et al., 2015). Less is known about the alpha oscillatory dynamics underlying the WM encoding period, although posterior alpha power has been shown to decrease in this stage, likely reflecting attentional processes (Heinrichs-Graham and Wilson, 2015). Likewise, alpha frequency has been linked to WM performance as a trait variable at rest (Klimesch, 1999) and during task performance in younger adults (Haegens et al., 2014).

Much of the research investigating age and load-related changes in alpha activity during WM have involved lateralised, visual WM tasks. In a study involving a hemifield change detection task with spatial cueing, it was found that while younger adults demonstrated higher alpha power ipsilateral to the attended hemifield during the retention period at medium and high loads, older adults only showed lateralisation at medium loads. This suggests that inhibitory processes indexed by alpha power modulation are not present in older adults when task difficulty increases (Sander et al., 2012). Likewise, a study employing a hemifield change detection task with an alerting cue showed that older adults had reduced alpha amplitude lateralisation during the retention period compared with younger adults, though between-load differences in alpha lateralisation did not predict performance at high WM loads in each age group (Tran et al., 2016). Lastly, using a lateralised delay match-to-sample paradigm with spatial cueing, it was found that at matched WM difficulty, alpha lateralisation during retention was minimal in older adults due to bilateral reductions in alpha power, while younger adults demonstrated lower alpha power contralateral to the attended hemifield (Leenders et al., 2018).

However, as the aforementioned studies involved visual WM tasks and a lateralised approach, and the majority of prior work investigating alpha activity during WM using Sternberg tasks have only included younger adults, it is unclear whether alpha activity contributes to verbal WM performance in older adults. A recent study employing magnetoencephalography during a high load (6-letter) modified Sternberg task reported that increases in visual alpha power during the WM maintenance period were present in both older and younger adults (Proskovec et al., 2016). However, relative to younger subjects, the increase in alpha activity was more rapid, widespread and persistent for longer in older adults, which was interpreted to reflect a compensatory mechanism to aid WM performance in older age (Proskovec et al., 2016). However, as WM load was not manipulated in this study, it is unclear whether older adults modulate visual alpha activity in order to facilitate verbal WM performance under varying WM loads. Likewise, while previous studies have found evidence for task- and load-related alpha frequency modulation during WM, these studies have only involved younger adults (Babu Henry Samuel et al., 2018; Haegens et al., 2014).

In the present study we investigated the age-related differences in visual alpha activity during verbal WM in response to increasing memory load. We applied a modified Sternberg task with 1-letter, 3-letter and 5-letter load conditions where WM processes were temporally delineated, in order to identify the alpha oscillatory dynamics underlying the WM encoding and retention stages. We ensured that any observed changes in the power of alpha oscillations were not due to age-related changes in peak alpha frequency by matching power measurements to individual alpha peaks. We sought to test the following hypotheses. First, older adults will show greater performance deficits at higher WM loads than younger adults. Second, older adults will show increased load-dependent modulation of visual alpha power during WM encoding and retention compared to younger adults. Third, age-related differences in visual alpha power during WM will correlate with task performance. Finally, cognitive reserve refers to the ability to maintain cognitive function in the presence of age-related changes to the brain, and can be acquired through socially and cognitively enriching activities throughout the lifetime (Barulli and Stern, 2013). As a secondary aim, we investigated whether cognitive reserve in older adults was associated with WM performance and alpha activity during the task.

## 2 Method

### 2.1 Participants, Demographics and Cognitive Reserve

24 younger adults (mean age: 23.2 years, SD: 4.60, range: 18-35 years, 8 male) and 30 older adults (mean age: 62.7 years, SD: 9.09, range: 50-86 years, 17 male) participated in the study. The samples in each group were not significantly different for years of education (older adults: M=15.87 years, SD=4.45 years; younger adults: M=15.71 years, SD=1.97 years, *t*_43.51_=0.182, *p*=0.857). All older adults were without cognitive impairment (Addenbrooke’s Cognitive Examination score (ACE-III) >82) (Mioshi et al., 2006). Cognitive reserve (CR) was calculated for each older adult participant by z-transforming their total years of education and scores on the National Adult Reading test (NART), which is used to estimate verbal IQ (Blair and Spreen, 1989). The z-scores were then averaged to form a cognitive reserve score, and participants were divided into high and low cognitive reserve groups using a median split. Exclusion criteria were a history of neurological or psychiatric disease, use of central nervous system altering medications, history of alcohol/substance abuse, uncorrected hearing/visual impairment and an ACE-III score of less than 82. All participants gave informed written consent before the commencement of the study, and the experiment was approved by the University of Adelaide Human Research Ethics Committee.

### 2.2 Working memory task

The modified Sternberg WM task used stimuli presented by PsychoPy software (Peirce, 2007) (figure 1). At the beginning of each trial, the participant fixated on a cross in the centre of the screen for 2 s. A memory set consisting of either 1,3 or 5 consonants was then shown for 1 s, followed by a 4 s retention period. For load-1 and load-3 trials, the consonants were presented centrally, with filler symbols (#’s) added to maintain equal sensory input for each condition. A probe letter was then shown, and the subject was instructed to press the right arrow key on a standard keyboard if the letter was in the memory set, or the left arrow key if it was not. The probe remained on the screen until the subject responded. Probe letters were present in the memory set at 50% probability. Participants received a practice block of 20 trials to familiarise themselves with the task, before performing 20 blocks of 15 trials, yielding 300 trials overall (i.e. 100 trials per load). Each block contained an equal number of trials for each load, presented pseudorandomly and a short break was allowed between blocks.

**Figure 1.**
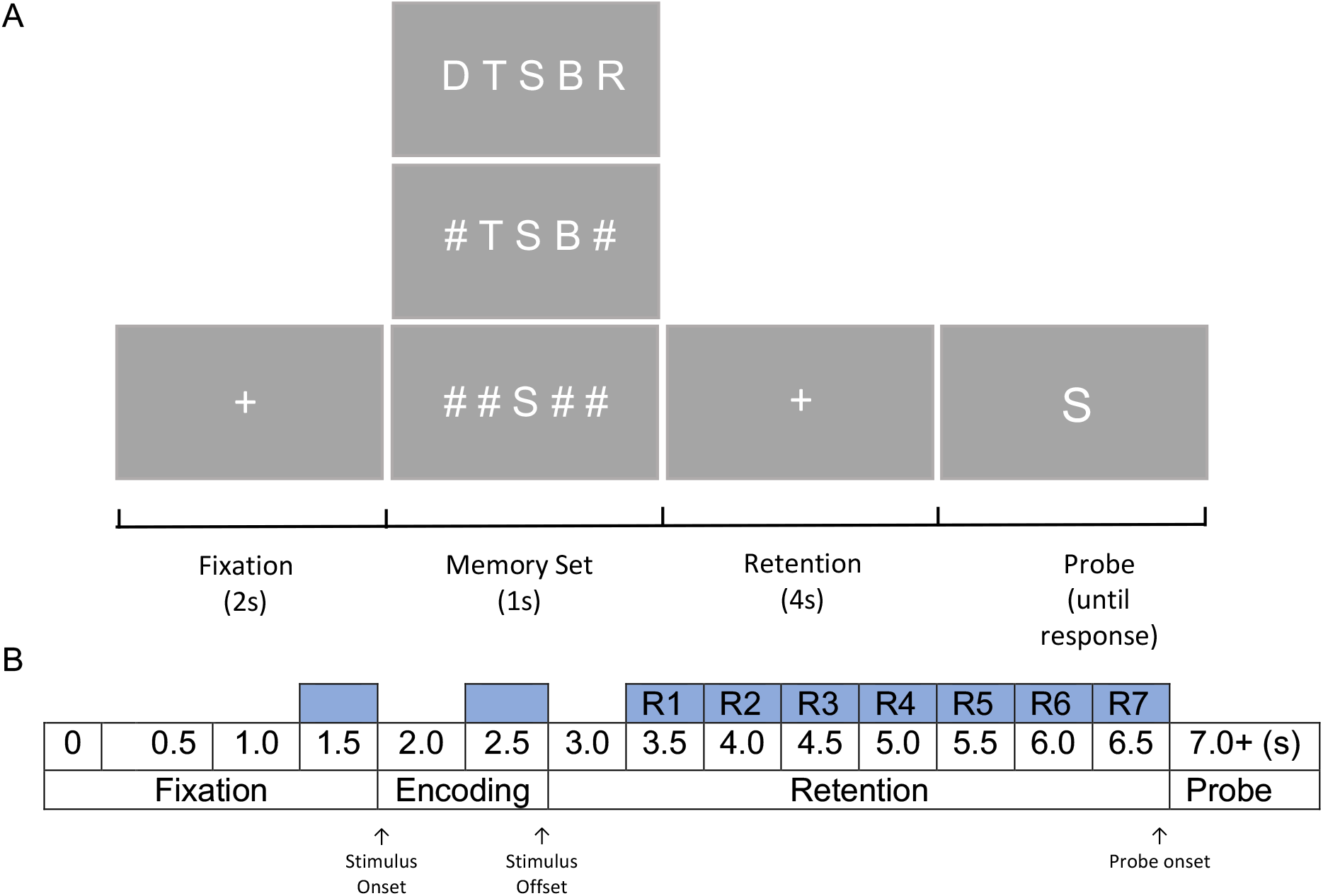
(A) Modified Sternberg WM task. Each trial contained four stages, including fixation lasting for 2 s; encoding, where a 1, 3 or 5 load memory set was displayed for 1 s; a 4 s retention stage and a retrieval stage where the subject responded to whether the probe was part of the memory set. (B) Schematic for EEG analysis periods (shown in blue) for fixation, encoding and retention.

To quantify WM performance, both accuracy (% correct) and reaction time (RT) for correct trials were calculated for each load condition.

### 2.3 EEG Data Acquisition

EEG data were recorded from 62 Ag/AgCl scalp electrodes arranged in a 10-10 layout (Waveguard, ANT Neuro, Enschede, The Netherlands) using a Polybench TMSi EEG system (Twente Medical Systems International B.V, Oldenzaal, The Netherlands). Conductive gel was inserted into each electrode using a blunt-needle syringe in order to reduce impedance to <5 kΩ. The ground electrode was located at AFz. Signals were amplified 20x, online filtered (DC-553 Hz), sampled at 2048 Hz and referenced to the average of all electrodes. EEG was recorded during each block of 15 trials of the WM task.

### 2.4 Data Pre-processing

Task EEG data were pre-processed using EEGLAB (Delorme and Makeig, 2004) and custom scripts using MATLAB (R2018b, The Mathworks, USA). Each block of EEG data was merged and incorrect trials, as well as trials with outlier RT (defined as >3xSD) were flagged for removal at the epoch stage.

Noisy and unused channels were then removed based on visual inspection, with an average of 2 channels removed from each age group (range old: 1-5, range young: 1-7). The data were then band-pass (1-100 Hz) and band-stop (48-52 Hz) filtered using zero-phase fourth-order Butterworth filters, down-sampled to 256 Hz and epoched −6s to 1s relative to the beginning of the probe. Only correct trials were included in further analysis. Independent component analysis (ICA) was then conducted using the FastICA algorithm (Hyvärinen and Oja, 2000), with the ‘symmetric approach’ and ‘tanh’ contrast function to remove artefacts resulting from eye-blinks and persistent scalp muscle activity. Data were then checked for remaining artefact via visual inspection and trials were removed if necessary (e.g. remaining blinks, non-stereotypic artefacts). Remaining trials were then split according to memory load condition.

After removing trials due to incorrect answers, outlier RTs or excessive artefact, on average, 254 trials were accepted in the final analysis for younger adults (range 171-294). 85 trials were retained in load-1 and load-3 and 84 in load-5. An average of 270 trials were accepted for final analysis for older adults (range 198-297). 273 trials were retained in load-1 and load-3, and 267 in load-5. A mixed effects linear model revealed a significant difference in the number of remaining trials across age groups *F*_1,52_ = 9.0, *p* = 0.04, but not across WM loads *F*_2,104_ = 2.8, *p* = 0.07, nor an age by load interaction, *F*_2,104_ = 0.7, *p* = 0.5.

### 2.5 Spectral Analysis

FieldTrip toolbox (Oostenveld et al., 2011) was used to analyse task EEG data. Time frequency representations of power to a 0.5 Hz frequency resolution were performed using a multi-taper time-frequency transformation based on multiplication in the frequency domain, a time window 3 cycles long and a Hanning taper. Power was calculated for individual trials before averaging for each load condition. The first 0.5 s of the encoding and retention periods were excluded to avoid spectral contributions from stimulus evoked responses to the memory set (figure 1B) (Babu Henry Samuel et al., 2018; Wang and Ding, 2011).

To account for age-related slowing of alpha (Klimesch, 1999), the alpha band frequency range was defined for each participant based on their peak alpha frequency at each stage of the task (fixation, encoding, retention) and for each load. Alpha frequency range was defined as 2 Hz above and below the peak frequency between 6 to 13 Hz (Klimesch, 1999). Alpha power was then averaged over this frequency range and across parieto-occipital and occipital electrodes (PO7, PO5, PO3, POz, PO4, PO6, PO8, O1, Oz and O2) at each WM stage (fixation, enocding and retention; figure 1B), as well as during each 0.5 s segment of the retention period.

### 2.6 Statistical Analyses

Statistical analyses were performed using R version 3.4.2. Mixed effects linear models were used to analyse the behavioural and neurophysiological data. For behavioural data, performance (RT and accuracy) was the outcome variable, WM load and age were fixed effects and subjects as the random effect. To investigate whether cognitive reserve influenced behavioural performance in the older adult group, a mixed effects model was conducted with fixed effects of load and cognitive reserve group, RT or accuracy as the outcome variable and subject as the random effect. For neurophysiological data, alpha power and alpha frequency were the outcome variable, age, WM load, WM stage and cognitive reserve (older adults only) were fixed effects and subjects as the random effect. Alpha power was log-transformed to normalise the data. Post-hoc pairwise t-tests were performed in case of significant main effects or interactions, with Bonferroni correction for multiple comparisons. Associations between alpha power (calculated as a change from fixation) and task performance were performed using Spearman’s correlation. In all tests, a p-value of less than 0.05 was considered statistically significant. Data were presented as mean ± SD in text and mean ± SEM in figures.

## 3 Results

### 3.1 Behavioural Results

While all participants performed the task successfully, task performance differed between memory load and age groups. A mixed effects linear model revealed significant main effects of age (*F*_1,52_ = 47.5, *p*<0.001) and load (*F*_2,104_= 241.3, *p*<0.001) on RT, with a significant age by load interaction (*F*_2,104_ = 17.8, *p*<0.001). Bonferroni corrected post-hoc tests revealed that younger adults responded significantly faster than older adults on load-1, load-3 and load-5 trials (*p*<0.001 for all). Likewise, RT for load-5 trials was significantly slower than load-3 and load-1 trials, and load-3 was significantly slower than load-1 in both age groups (*p*<0.001 for all) (figure 2A).

**Figure 2.**
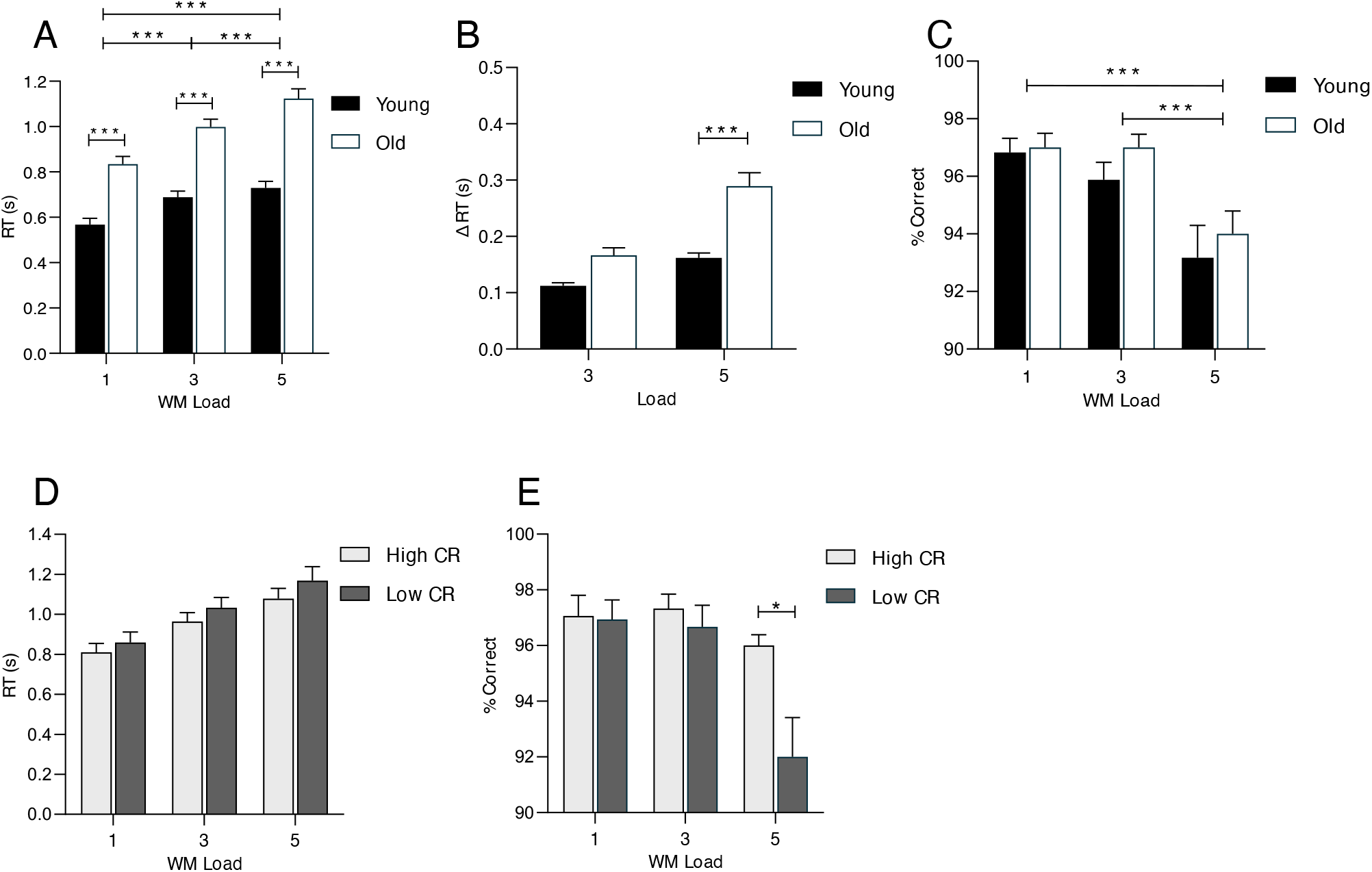
(A) RT for correct responses, (B) change in RT from load-1, and (C) percentage of correct responses for each WM load in younger and older adults. (D) RT for correct responses and (E) percentage of correct responses for older adults with high and low cognitive reserve. * *p* <0.05 \*\*\**p*<0.001.

To examine the interaction between age and load on RT, we examined the change in RT relative to load-1 between age-groups. A mixed effects linear model revealed significant main effects of age (*F*_1,52_ = 18.97, *p*<0.001) and load (*F*_2,52_ = 94.96, *p*<0.001) on RT, with a significant age by load interaction (*F*_2,52_ = 15.88, *p*<0.001). Bonferroni corrected post-hoc tests revealed that the increase in RT from load-1 to load-5 was larger in older adults compared to younger adults (*p*<0.001). The increase in RT from load-1 to load-3 did not differ between age groups (*p*=0.16) (figure 2B).

The model revealed a significant main effect of load on accuracy (*F*_2,104_= 19.7, *p*<0.001). There was no significant main effect of age (*F*_1,52_ = 0.91, *p* = 0.34), nor a significant interaction between age and load (*F*_1,104_= 0.4, *p* = 0.7). Post-hoc tests revealed that accuracy was significantly lower for load-5 trials compared to both load-1 (p<0.001) and load-3 trials (p<0.001) but did not differ between load-1 and load-3 trials (p=0.542) (figure 2C).

A composite score of education years and NART results was used to calculate cognitive reserve in the older adults. The average score on the NART was 38.7 ± 5.25, which corresponds to an average pre-morbid verbal IQ of 106.6 ± 6.8. There was no significant difference in the age of the low and high cognitive reserve groups (*t*_28_ = 0.27, *p*=0.79). For RT, the model revealed a significant main effect of load (*F*_2,56_ = 123.4, *p*<0.01), but no main effect of cognitive reserve (*F*_1, 28_ = 0.92, *p* = 0.34), nor a load by cognitive reserve interaction (*F*_2, 56_= 0.62, *p*=0.54). For accuracy, the model revealed a significant main effect of load (*F*_2, 56_= 12.51, *p* <0.001), a non-significant main effect of cognitive reserve (*F*_1, 28_= 3.59, *p*=0.07) and a significant load by cognitive reserve interaction (*F*_2,56_= 4.58, *p*=0.014). Bonferroni corrected post-hoc tests revealed that in load-5 trials, older adults with higher cognitive reserve performed better than those with low cognitive reserve (*p*=0.028). No significant differences in accuracy were found between the high and low cognitive reserve group in load-3 (*p*=0.99) or load-1 (*p*=0.99) trials.

Given that prior studies have demonstrated sex differences in verbal WM performance at high loads (Reed et al., 2017), we examined whether sex differences in RT and accuracy were present in our sample. A mixed effects linear model revealed no main effect of sex on RT (*F*_1, 50_ = 0.49, *p*=0.48), nor interactions between sex and age (*F*_1, 50_ = 0.11 *p*=0.74), sex and load (*F*_2,100_ = 0.35, *p*=0.70) or age, sex and load (*F*_2,100_ = 3.1, *p*=0.051). Likewise, there was no main effect of sex on accuracy (*F*_1,50_ = 1.2, *p*=0.28), nor interactions between sex and age (*F*_1, 50_ = 0.14, *p*=0.71), sex and load (*F*_2,100_ = 1.6, *p*=0.21), or age, sex and load (*F*_2, 100_ = 0.08, *p*=0.92).

### 3.2 Alpha frequency and power modulation

Power spectra for young and older adults at each load and WM stage are shown in figure 3. Participants in which an alpha peak was not detected at any WM stage or load were excluded from analysis of alpha peak frequency (4 older adults). A linear mixed effects model revealed significant main effects of age (*F*_1,48_ = 4.7, *p* = 0.04), WM stage (*F*_2,347_ = 8.0, *p* <0.001) and load (*F*_2,347_ = 5.8, *p* = 0.009). There were no significant interactions. Bonferroni corrected post-hoc tests revealed that on average, older adults had lower alpha frequency than younger adults (*p*=0.04) (figure 3B). Alpha frequency was significantly higher in load-5 compared with load-1 trials (*p*=0.008), but was not different between load-3 and load-5 trials or load-1 and load-3 trials (Figure 3C). Alpha frequency was significantly higher during encoding compared with fixation (p =0.006) and retention (p=0.003), but did not differ between retention and fixation (Figure 3D).

**Figure 3.**
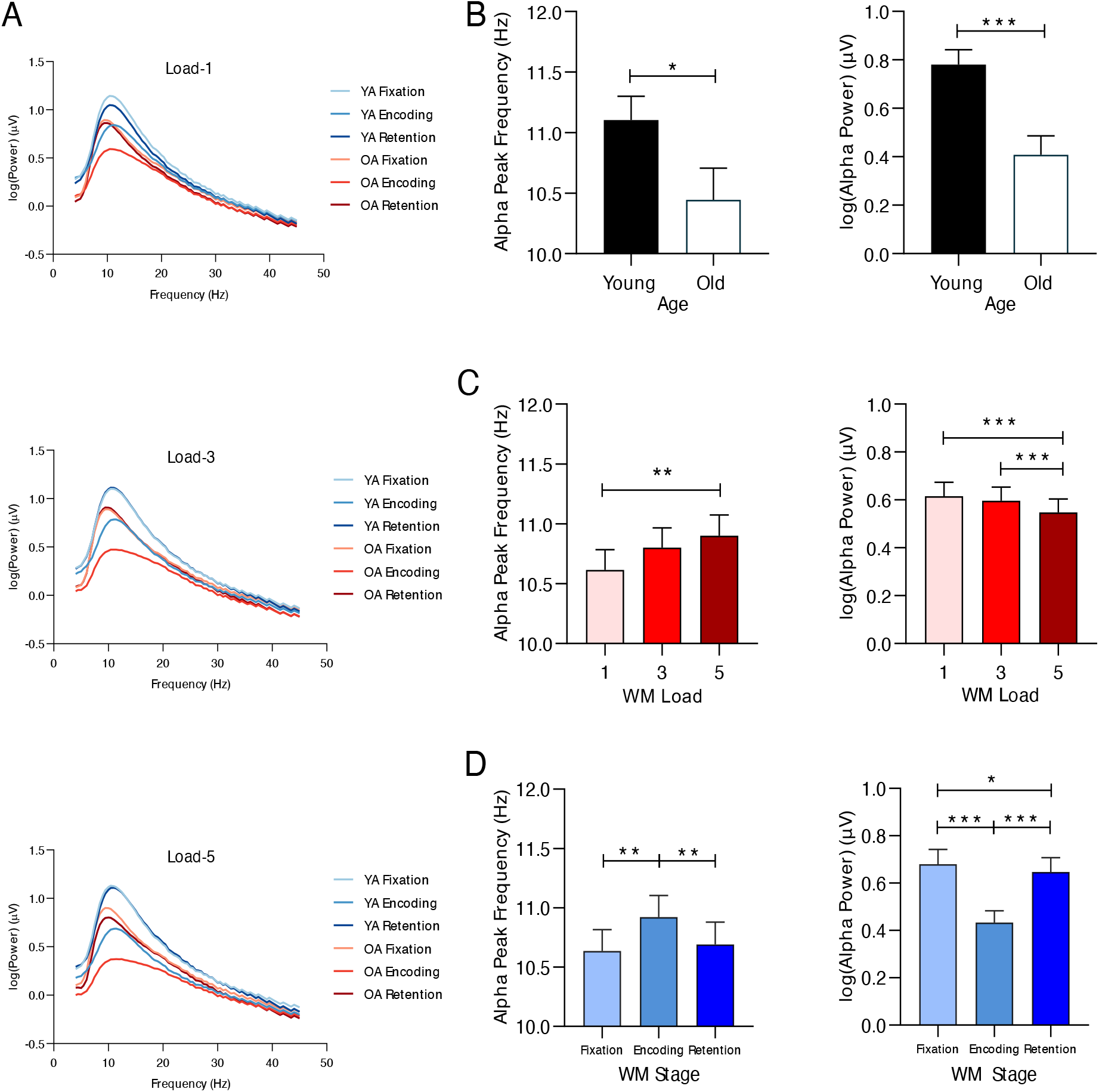
(A) Power spectra for young (YA) and older adults (OA) during each WM stage for load-1 (top), load-3 (middle) and load-5 (bottom) trials. (B-D) Peak alpha frequency (left) and alpha power (right) between (B) age groups, (C) WM load and (D) stages of the WM task. * p <0.05, ** p <0.01, *** p<0.001

Alpha power was calculated using individual peak frequency at each WM stage (fixation, encoding, retention) for each load. If a peak was not found in the retention period, the value for fixation was used to determine the frequency band for power calculations. If no peaks were found in any WM stage, the participant was excluded from further analysis (4 older adults). A mixed model revealed main effects of age (*F*_1,48_ = 13.5, *p*<0.001), WM stage (*F*_2,384_ = 240.4, *p*<0.001) and load (*F*_2,384_ = 16.6, *p*<0.001), as well as an age x WM stage (*F*_2,384_ = 3.6, *p*=0.03) and WM stage x load (*F*_4,384_ = 7.1, *p*<0.001) interaction. Bonferroni corrected post-hoc tests revealed that overall, alpha power was significantly lower in older adults compared with younger adults (*p*<0.001) (figure 3B). Alpha power was significantly higher in load-1 trials compared with load-5 trials (p<0.001), in load-3 trials compared with load-5 trials (p<0.001), but not different between load-1 and load-3 trials (figure 3C). Alpha power was significantly lower during encoding compared with both fixation (*p*<0.001) and retention (*p*<0.001), and alpha power during retention was significantly lower than in fixation (p=0.03) (figure 3D).

To examine the interaction between age and WM stage, mixed models were conducted separately in each age group with alpha power as the outcome variable, WM stage as the fixed effect and subject as the random effect. In older adults the model was significant (*F*_2, 206_=105.3, p <0.001), with Bonferroni corrected post-hoc tests revealing that for older adults, alpha power was significantly lower during encoding compared with fixation (p<0.001) and retention (p<0.001), and that alpha power during retention was significantly lower than during fixation (p=0.003). The model was also significant in younger adults (*F*_2, 190_=108.8, p <0.001), with Bonferroni corrected post-hoc tests revealing that for younger adults, alpha power during encoding was lower compared with both fixation (p<0.001) and retention (p<0.001), but there was no difference between fixation and retention (figure 5A). However, an independent samples t-test revealed that the change in alpha power from fixation to retention was not significantly different between age groups (*t*_42_=-1.4, *p*=0.17).

To investigate the interaction between WM stage and load, a mixed model was conducted for each WM stage, with load as the fixed effect and subjects as the random effect. For alpha power during the fixation period, the model was significant (*F*_2,98_ = 3.4, *p* = 0.004), though Bonferroni corrected post-hoc tests revealed no differences in alpha power between loads. For alpha power during encoding, the model was significant (*F*_2,98_ = 66.3, *p*<0.001). Bonferroni corrected post-hoc tests revealed that during encoding, alpha power decreased with increasing memory load (all comparisons p<0.001). For alpha power during retention, the model was significant (*F*_2,98_ = 11.7, *p*<0.001). Bonferroni corrected post-hoc tests revealed that during retention, alpha power was significantly lower in load-5 trials compared with both load-1 (*p*=0.002) and load-3 trials (*p*<0.001), but did not differ between load-1 and load-3 trials (figure 4B).

**Figure 4.**
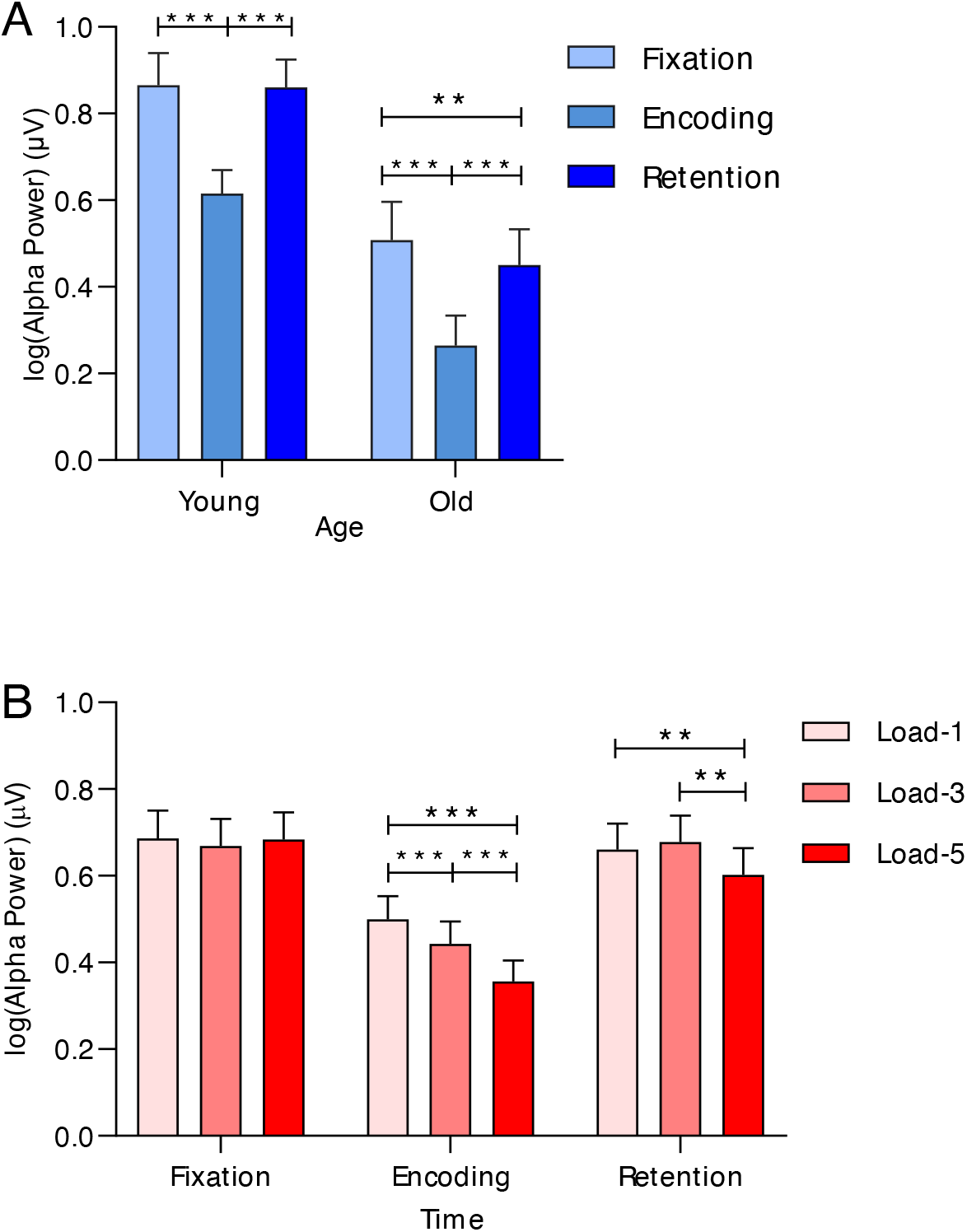
Alpha power modulation during the stages of the WM task between age groups (A) and WM loads (B) * *p* <0.05, ** *p* <0.01, *** *p*<0.001

When examining the effect of cognitive reserve group on alpha activity in older adults, the model for alpha frequency revealed no significant main of WM stage (*F*_2, 157_ = 1.71, *p* = 0.18), cognitive reserve group (*F*_1, 24_=2.07, *p*=0.16) or load (*F*_2, 15_=2.36, *p*=0.097), and no significant interactions. For alpha power, the model revealed main effects of WM stage (*F*_2, 192_ = 118.6, *p* < 0.001), load (*F*_2, 192_ = 10.7, *p* < 0.001) and a WM stage by load interaction (*F*_4, 192_ = 3.75, *p* = 0.006), but no main effects or interactions involving cognitive reserve (*p* ≥ 0.41 for all).

### 3.3 Retention period time course

For closer inspection of the temporal changes during the retention period, we calculated alpha power and peak frequency for each 0.5s segment of the retention period. Only participants who had an alpha peak at each time point during the retention period were included in this analysis (19 older adults, 23 younger adults).

A mixed model with peak alpha frequency as the outcome, age, load and time as fixed effects, and subjects as the random effect revealed main effects of time (*F*_7,982_ = 20.2, *p*<0.001) and load (*F*_2,982_ = 23.0, *p*<0.001). There were no other significant main effects or interactions. Bonferroni corrected post-hoc tests revealed that in R1 (i.e. 0.5-1s from the start of the retention period), alpha frequency was higher than in all subsequent time increments (all *p*<0.001). Likewise, alpha frequency was higher in load-5 during retention when compared with load-1 (*p*<0.001) and load-3 (*p*<0.001) (figure 5A).

**Figure 5.**
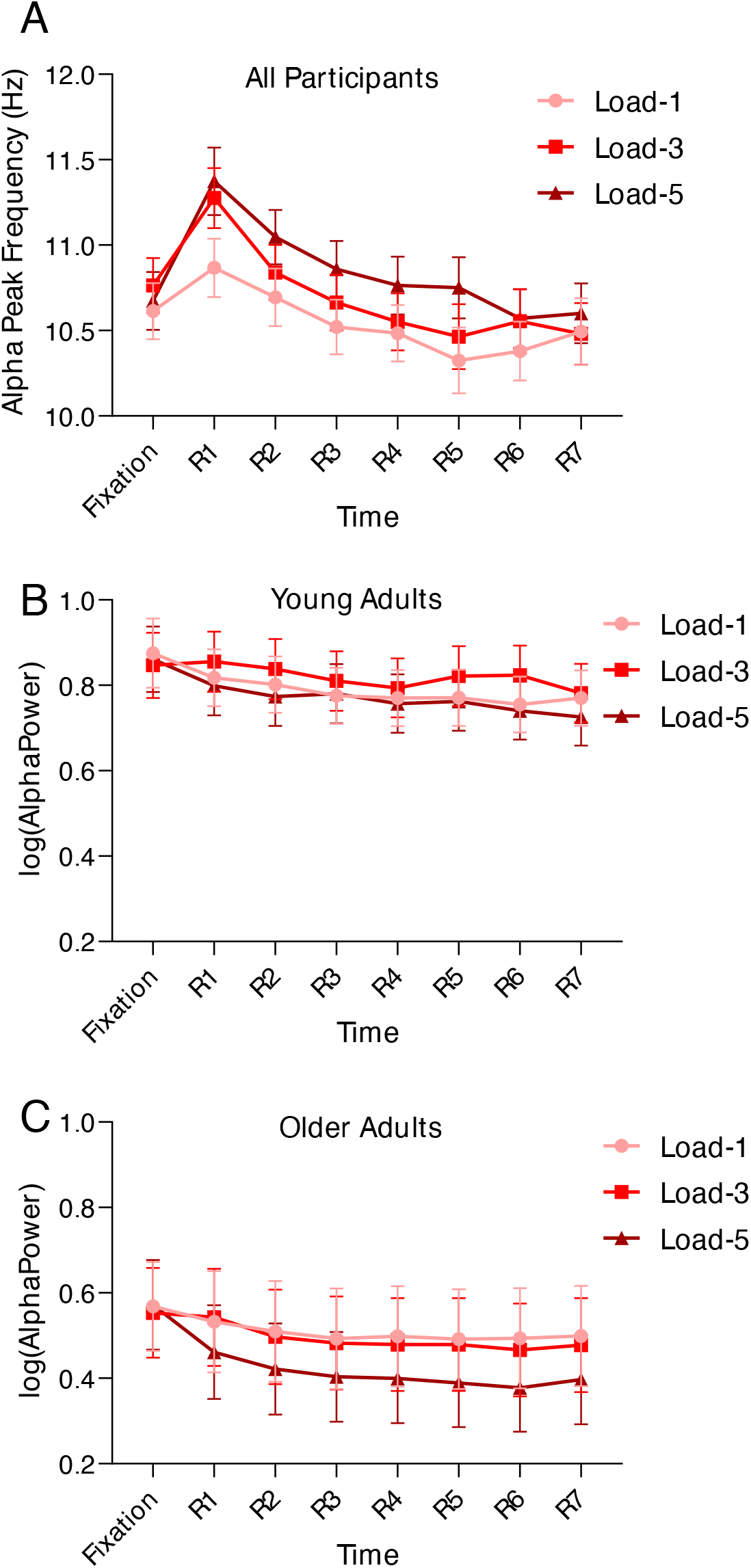
(A) Alpha peak frequency modulation across time and load during fixation and retention for all participants. (B-C) Alpha power modulation across time and load during fixation and retention for (B) young and (C) older adults.

A mixed model with alpha power as the outcome, age, load and time as fixed effects, and subjects as the random effect revealed main effects of age (*F*_1,440_ = 109.4, *p*=0.025), time (*F*_3,440_ = 16.7, *p*<0.001) and load (*F*_2,440_ = 15.5, *p*<0.001). There were no significant interactions. Bonferroni corrected post-hoc tests revealed that older adults had lower alpha power during fixation and retention than younger adults (*p*=0.007). Alpha power was significantly greater in the fixation period compared with each time point except R1. Further, alpha power was lower in load-5 trials compared with load-1 (*p*<0.001) and load-3 (*p*<0.001), but there were no differences between load-1 and load-3 (figure 5B).

### 3.4 Relationship between alpha power and task performance

Spearman correlation analyses revealed no significant association between alpha power during the encoding or retention period (relative to fixation) and performance metrics for all WM loads in both age groups (table 1).

**Table 1.** Coefficients for correlations between RT and accuracy and alpha power during encoding and retention, relative to fixation, at all WM loads for younger and older adults.

|  |  | Encoding |  |  |  | Retention |  |  |  |
| --- | --- | --- | --- | --- | --- | --- | --- | --- | --- |
|  |  | Young |  | Old |  | Young |  | Old |  |
|  |  | RT | Accuracy | RT | Accuracy | RT | Accuracy | RT | Accuracy |
| Load-1 | *rho* | -0.17 | 0.16 | 0.10 | -0.20 | -0.16 | 0.01 | 0.02 | -0.22 |
|  | *p* | 0.43 | 0.46 | 0.62 | 0.34 | 0.45 | 0.95 | 0.94 | 0.28 |
| Load-3 | *rho* | -0.21 | 0.26 | -0.09 | -0.12 | -0.20 | -0.01 | 0.01 | -0.15 |
|  | *p* | 0.33 | 0.23 | 0.68 | 0.56 | 0.34 | 0.65 | 0.97 | 0.47 |
| Load-5 | *rho* | -0.28 | 0.13 | -0.06 | 0.16 | -0.22 | -0.23 | -0.02 | -0.1 |
|  | *p* | 0.18 | 0.54 | 0.75 | 0.43 | 0.31 | 0.29 | 0.92 | 0.77 |

To further examine whether alpha frequency and power influenced WM performance, we sorted trials within each subject to a high or low RT group according to a median split. Alpha frequency and power were then averaged over each RT group for each load during the encoding and retention periods. A mixed model with alpha frequency as the outcome, age, load, WM stage and RT group as fixed effects and subject as the random effect revealed a main effect of age (*F*_1,50_ = 12.37, *p*<0.001), but no main effects or interactions involving RT group (*p*≥0.09 for all). Likewise, a mixed model with alpha power as the outcome revealed main effects of WM stage (*F*_1,572_ = 646.2, *p*<0.001), age (*F*_1,52_ = 13.77, *p*<0.005) and load (*F*_2, 572_ = 26.26, *p*<0.001), but no main effects or interactions involving RT group (*p*≥0.08 for all).

## 4 Discussion

In this study, we investigated age-related differences in visual alpha power and frequency during the encoding and retention stages of WM in response to varying loads. Behaviourally, older adults were slower to respond at all WM loads compared to younger adults, but there were no age differences in accuracy. However, older adults with higher cognitive reserve performed better on load-5 trials compared with those with lower cognitive reserve. Overall, both alpha frequency and power were lower in older adults than in younger adults in each WM stage. During encoding, alpha power decreased with increasing WM load and alpha frequency increased. Regardless of age, alpha power was lower in load-5 than in load-1 and load-3 trials, but alpha frequency increased with load during retention. While alpha power during retention was lower than fixation in older, but not younger adults, the relative change from fixation was not significantly different between age groups. Further, individual differences in visual alpha power did not predict individual task performance within age groups, at any WM loads.

### 4.1 At all WM loads, older adults are slower to respond to the probe than younger adults, but accuracy does not differ across age groups

As expected, older adults were slower to respond to the probe at all WM loads compared with younger adults. However, we found no age differences in accuracy, with many participants from both age groups performing at near ceiling level for accuracy. Therefore, the difference in RT in the older group likely does not reflect WM deficits, but rather age-related changes in processing speed (Salthouse, 1996). It has been shown that verbal WM might be more resistant to age effects than visual WM (Hale et al., 2011), and it is possible that our task was not difficult enough to capture age-differences. Taken together, we did not find strong evidence for working memory impairment with ageing in our sample, although this may be specific to the type of task performed.

### 4.2 Alpha power is modulated by load during the encoding and retention period for both younger and older adults

Alpha suppression occurred during the encoding period in both age groups, with a strengthening of this response with increasing WM load. Alpha suppression has long been thought to reflect attentional processes (Klimesch, 1997), as when attention is directed to external visual events, alpha power in visual cortex decreases with increasing attention demands (Rajagovindan and Ding, 2010; Sauseng et al., 2005). Therefore, a decrease in alpha power during encoding likely reflects an increase in cortical excitability to enhance stimulus processing (Heinrichs-Graham and Wilson, 2015; Klimesch, 1997; Murphy et al., 2019; Romei et al., 2010; Thut et al., 2011). Although alpha power was lower in older relative to younger adults, our results suggest that alpha suppression during encoding follows a similar pattern across age groups. This is consistent with previous studies that have shown that suppression processes during the encoding period, as indicated by alpha activity, remain relatively intact in older adults (Gazzaley et al., 2008). However, even though both age groups demonstrated poorer performance with increasing WM load, individual differences in alpha suppression during encoding did not support WM performance under varying loads in our task.

In both age groups, we found that alpha power decreased under higher WM loads during the retention period. This contrasts with the previously reported increase in visual alpha power during retention in younger adults completing modified Sternberg tasks (Jensen et al., 2002; Proskovec et al., 2019; Tuladhar et al., 2007; Wang et al., 2016). Though an increase in alpha power during retention has been interpreted to reflect inhibition of task irrelevant information, in lateralised tasks, alpha power decreases in task-relevant brain regions, but increases in task-irrelevant regions, and the magnitude of this reduction correlates with WM load (Sauseng et al., 2009). Likewise, in a study employing a delayed match-to-sample task, stronger alpha suppression during the retention period was seen under higher visual WM loads (Fukuda et al., 2015). Despite our use of verbal rather than visual stimuli, given that the magnitude of alpha suppression was greatest in load-5 during retention, the pattern of alpha activity we observed may reflect a mechanism for holding multiple items in WM rather than inhibition of task-irrelevant brain regions.

In terms of age-related findings, our results contrast with a recent study employing a 6-letter modified Sternberg task, where it was observed that older adults exhibited a greater increase in visual alpha power during the retention period compared to younger adults (Proskovec et al., 2016). This was interpreted in that study to align with the Compensation-Related Utilisation of Neural Circuits Hypothesis (CRUNCH) (Reuter-Lorenz and Cappell, 2008), which suggests that generally, people recruit more brain regions when task-difficulty increases. Older adults are thought to recruit more cortical regions at lower loads than younger adults to compensate for cognitive decline. In our study, however, while younger adults demonstrated no difference in alpha power during the retention period compared with fixation, older adults demonstrated a decrease in power from fixation, regardless of load. While this difference in alpha power relative to fixation was not significantly different between age groups, it may not be physiologically feasible for older adults to modulate visual alpha power in a range that is behaviourally advantageous during WM due to lower resting alpha power. Conversely, if alpha suppression is indicative of the active maintenance of WM representations, the decrease in alpha power seen at load-5 in older adults may be another form of compensatory neural strategy. As such, clarifying the role of alpha suppression during WM and cognitive ageing is a topic for future research.

Further, studies investigating the alpha rhythm in both younger and older adults tend to define alpha as a narrow band (usually 8-12Hz) and average over spectral activity in that range for all subjects. Given that peak alpha frequency decreases with age, alpha power may fall outside of the fixed alpha frequency band, or activity in theta/beta frequencies may be included in the alpha window. Our results show that when alpha power is calculated based upon individual peak alpha frequency, the pattern of alpha activity across WM stages in older adults appears similar to that of younger adults but lower in magnitude, even when WM performance is matched to that of younger adults.

### 4.3 Age, task and load modulation of alpha frequency

Age has long been known as one of the most important factors influencing the frequency of the alpha rhythm (Dustman et al., 1985; Klimesch, 1997). Resting state alpha peak frequency has been shown to be a stable neurophysiological trait in healthy younger and older adults (Grandy et al., 2013), however, it is becoming increasingly clear that alpha peak frequency shifts during cognitive tasks. In particular, a study employing an n-back task demonstrated a load-dependent increase in alpha frequency in healthy young adults (Haegens et al., 2014), while in a modified Sternberg task, a load-dependent decrease in alpha frequency during encoding and an increase during retention were apparent (Babu Henry Samuel et al., 2018).

During the encoding and retention periods, we observed a load-dependent increase in peak alpha frequency, suggesting that alpha frequency reflects cognitive engagement or is a metric of cognitive load that is common to both younger and older adults. Considering alpha activity is associated with inhibitory processes, slower alpha frequency would allow for longer windows of suppression (Jensen and Mazaheri, 2010; Sadaghiani and Kleinschmidt, 2016), which may facilitate protection against interference during WM. Our results only partially support this idea. According to this interpretation, the increase in alpha frequency seen during encoding may reflect a release of inhibition to facilitate information processing; consistent with the decrease in alpha power observed during encoding in this study. However, the fact we also observed load-dependent increases in frequency during retention may invalidate this, as higher alpha frequency during retention should be counterproductive to performance. This was shown in a recent study which demonstrated that higher peak frequency during retention led to slower RT (Babu Henry Samuel et al., 2018). Therefore, determining the task-relevance of alpha peak frequency during WM is a topic for future research.

### 4.4 Cognitive Reserve

Within the older adult group, we found that participants with higher composite cognitive reserve performed more accurately on the WM task at load-5, but not at load-3 and load-1, than those with low cognitive reserve. A previous study employing a verbal WM Sternberg task that increased in load from 1 to 7 reported similar results, with subjects with higher cognitive reserve performing more accurately in the task at higher loads than those with lower cognitive reserve (Speer and Soldan, 2015). However, within the older adult group, we did not see differences in alpha power or frequency during the task between cognitive reserve groups. Theoretically, it is proposed that cognitive reserve does not directly alter age-related neural changes, but rather modifies the behavioural outcome of these anatomical or physiological changes (Barulli and Stern, 2013). However, cognitive reserve may influence other oscillatory dynamics at rest and during task performance that were not investigated in this study, presenting an avenue for further investigation.

### 4.5 Limitations

There are several limitations of this study. First, the age range of the older adult sample is much larger (36 years) than the younger adult sample (17 years). Given that trajectories of change in cognitive performance are largely heterogeneous across older adults (Hayden et al., 2011), future work should take into account individual differences in age-related WM decline. Second, though the modified Sternberg task used in this study allowed the temporal delineation of the encoding and retention stages of WM, this task does not assess the manipulation of items in WM (Baddeley, 1992). As such, further research is required to expand upon whether alpha activity is altered with age during the manipulation component of WM processing. Finally, lower alpha power seen with age might be due to structural brain differences such as atrophy in cortical tissue seen with age, brain size or skull thickness which are not able to be accounted for or assessed with EEG (Frodl et al., 2001). Future work may investigate how structural brain changes influence oscillatory activity recorded during WM task performance.

## Author Notes

NCR and MRG are supported by Australian Research Council Discovery Early Career Researcher Awards (180100741 and 200100575, respectively). SS is supported by an Australian Government Research Training Program (RTP) Scholarship. We would also like to thank the participants who dedicated their time to be involved in this study.

The authors confirm that there are no known conflicts of interest associated with this publication

